# The functional small RNA interactome reveals targets for the vancomycin-responsive sRNA RsaOI in vancomycin tolerant *Staphylococcus aureus*

**DOI:** 10.1101/2023.08.23.554412

**Authors:** Winton Wu, Chi Nam Ignatius Pang, Daniel Mediati, Jai Justin Tree

## Abstract

RNA-RNA interactome profiling techniques have expanded our understanding of sRNA-mRNA interactions in bacteria. However, determining the function of these interactions for hundreds of sRNA-mRNA pairs is a major challenge. At steady-state, protein and mRNA abundances are often highly correlated and lower than expected protein abundance may indicate translational repression of an mRNA. To specifically identify sRNA-mRNA interactions that regulate mRNA translation, we examined the correlation between gene transcript abundance, ribosome occupancy, and protein levels. We used SOMS to cluster genes with similar transcription and translation patterns and identified a cluster of mRNAs that appeared to be post-transcriptionally repressed. By integrating this clustering analysis with sRNA-mRNA interactome data generated in vancomycin tolerant *S. aureus* by RNase III-CLASH, we identified sRNAs that may be mediating this translational repression. We have confirmed sRNA-dependant post-transcriptional repression of several mRNAs in this cluster. Two of these interactions are mediated by RsaOI, a sRNA that is highly upregulated by vancomycin treatment. While RsaOI is not essential for vancomycin tolerance, we demonstrate regulation of the phosphocarrier protein HPr and the cell-wall autolysin Atl. These findings suggest that RsaOI may serve as a regulator of carbon metabolism and cell wall turnover during cell wall stress exerted by vancomycin.

## INTRODUCTION

The emergence of multidrug-resistant *Staphylococcus aureus* (MRSA) is a major public health concern. Current treatment is dependent on the efficacy of last line antibiotics like vancomycin. However, MRSA isolates that exhibit intermediate resistance (MIC 4-8 µg/mL) to vancomycin are increasingly detected worldwide and are associated with treatment failure (Howden *et al*, 2011). These vancomycin-intermediate *S. aureus* (VISA) isolates arise from the acquisition of a disparate series of point mutations that lead to physiological changes including cell wall thickening and reduced autolysis (Howden *et al*, 2010).

Transcriptional profiling has revealed that antibiotic treatment drives conserved changes in small RNA (sRNA) expression in *S. aureus* and may contribute to the VISA phenotype (Howden *et al*, 2013). Small RNAs have been established as global post-transcriptional regulators of bacterial gene expression which modulate mRNA expression by direct base-pairing. Small RNAs have been found to act by a wide array of mechanisms and have been shown to occlude the ribosomal binding site (Bouvier *et al*, 2008), activate translation (Frohlich & Vogel, 2009), recruit and occlude RNase recognition (Bandyra *et al*, 2012; Frohlich *et al*, 2013), control transcription termination (Sedlyarova *et al*, 2016), and modulate ribosome sliding to affect translational (Jagodnik *et al*, 2017). In addition, sponging and buffering interactions (Tree *et al*, 2014; Lalaouna *et al*, 2015, McKellar *et al*, 2022) with other non-coding RNAs expand individual sRNA networks across multiple regulatory nodes.

Recent progress using proximity-dependant ligation techniques such as CLASH, RIL-seq, and Hi-GRIL-seq have massively expanded the number of sRNA-mRNA interactions identified *in vivo* (Iosub *et al*, 2020; Matera *et al*, 2022; Melamed *et al*, 2016; Waters *et al*, 2017) (Mediati *et al*, 2022; Melamed *et al*., 2016; Waters *et al*., 2017). Using a technique termed CLASH (crosslinking, ligation, and sequencing of hybrids), sRNA-RNA interactions associated with the endoribonuclease RNase III were captured and sequenced in the methicillin resistant *S. aureus* (MRSA) strains JKD6009 and USA300 (Mediati *et al*, 2022; McKellar *et al*., 2022). This was the first *in vivo* map of RNA-RNA interactions in a Gram-positive bacterium and revealed a regulatory mRNA 3’ UTR that was required for vancomycin tolerance (Mediati *et al*., 2022). In parallel work, small RNA regulation of the alpha phenol soluble modulin toxins were uncovered (McKellar *et al*, 2022).

RNA-RNA interaction networks identify interacting RNA pairs but do not provide information on the function of the interaction. Follow-up studies on individual sRNA-mRNA interactions are required to determine the regulatory functions and biological significance of the interaction. These studies typically employee RNA-seq or proteomics to monitor changes in transcript or protein abundance after expression of a single sRNA and are not well suited to defining the functions of many different sRNAs on their cognate targets. These follow-up studies have also indicated that some sRNA-mRNA interactions recovered by *in vivo* proximity-dependant ligation do not affect mRNA transcript or protein abundance (Faigenbaum-Romm *et al*, 2020; Iosub *et al*., 2020). It is clear that for sRNA-RNA interaction networks recovered by proximity-dependant ligation, predicting the functional outcome of the interaction will require new approaches to determine systems-wide sRNA-mRNA functions.

Here we have captured the RNase III-associated RNA-RNA interactome of the VISA strain JKD6008. To identify mRNAs that are post-transcriptionally regulated we have analysed RNA-seq, Ribo-seq, and proteomics data to assess the correlation between mRNA abundance, translation, and protein abundance. Using this data, we have divided mRNAs into clusters using self-organising maps (SOMS) and overlaid these clusters with mRNAs interacting with sRNAs in our RNase III-CLASH network. This approach identified a cluster of mRNAs with below average correlation between mRNA and protein abundance that is enriched for sRNA-targeted mRNAs suggesting that sRNA regulation is responsible for translational silencing of these mRNAs. We confirm that this SOMS cluster contains translationally repressed sRNA targets by examining the vancomycin-induced sRNA, RsaOI. We identify functional targets for the sRNA, RsaOI, that are translationally repressed and promote mRNA turnover in our SOMS clusters. The RsaOI targetome includes the autolysin *atl* that is involved in cell wall turnover and provides a pathway for post-transcriptional control of cell wall autolysis in response to vancomycin treatment.

## MATERIALS AND METHODS

### Bacterial strains and culture conditions

Bacterial strains and plasmids used in this study are listed in Supplementary Table 1 while the oligonucleotides are listed in Supplementary Table 2. *E. coli* strains DH5α and IMO8B were routinely cultured at 37°C in Luria-Bertani (LB) media. To facilitate plasmid extraction in *E. coli*, LB media was supplemented with the following antibiotic concentrations when appropriate: 10 µg/mL chloramphenicol or 100 µg/mL ampicillin. On the other hand, *S. aureus* strains JKD6008 and JKD6009 were routinely cultured at 37°C in either Brain Heart Infusion media or Mueller-Hinton (MH) media. To facilitate plasmid selection in *S. aureus*, media was supplemented with the following antibiotic concentrations when appropriate: 10 µg/mL chloramphenicol and/or 10 µg/mL erythromycin. Construction of the RNase III-HTF tagged and deletion of RsaOI in *S. aureus* strain JKD6008 was performed through allelic exchange using the pIMAY-Z vector (Monk *et al*, 2012).

### RNase III-CLASH of vancomycin-intermediate *S. aureus* (VISA)

Preparation of the VISA RNase III-CLASH libraries was performed as outlined in our previous report (McKellar *et al*., 2022; Mediati *et al*., 2022) with some modifications. *S. aureus* JKD6008 wildtype, untreated JKD6008 RNase III-HTF and vancomycin treated (final concentration of 2 µg/mL for 10 minutes) JKD6008 RNase III-HTF cultures were grown in duplicates to an OD_600nm_ of 0.8 in MH media. Cultures were crosslinked with 400 mJ of UV-C (Vari-X-Link, UV03) and immediately harvested by vacuum filtration. Cells pellets were lysed and clarified by centrifugation. Clarified supernatants were added to pre-washed mouse-IgG agarose (A0919 Sigma-Aldrich) and incubated at 4°C for 1 hr to sequester the immunoglobulin-binding protein Sbi. The supernatant was transferred to pre-washed M2 anti-FLAG resin (Sigma-Aldrich) and incubated at 4°C for 1.5 hours. The resin was washed and incubated with 50 U of TEV protease and 100 µg/mL FLAG peptide (Sigma-aldrich) for 2 h at 18°C. The eluate was then collected and incubated with 0.15 U of RNace-IT (Agilent) at 20°C for 5 min. The eluate was added to pre-washed Pierce^TM^ Ni-NTA magnetic agarose slurry and incubated overnight at 4°C.

The RNase III.RNA complexes were washed and the 5′ ends of bound RNAs were dephosphorylated with the addition of thermosensitive alkaline phosphatase (Promega). RNA was radiolabelled by phosphorylation with T4 PNK (Sigma-aldrich) and ^32^P-γATP (PerkinElmer, cat no. BLU502A250UC) for 100 min at 20°C. The ^32^P-radioabelled RNase III.RNA complexes were washed and unique barcoded 5’ linkers and 3’ App-PE adapters were ligated to the bounded RNA using 40 U T4 RNA ligase I (NEB) for each ligation step. The ^32^P-radioabelled RNase III.RNA complexes were subsequently eluted in 60 µL of 1x NuPAGE LDS sample buffer (Sigma-aldrich) at 65°C for 10 mins and resolved on a NuPAGE 4–12% Bis-Tris PAGE gel (Invitrogen) at 120 V for 1 h. Complexes were visualized by autoradiography using the Typhoon^TM^ FLA 9500 gel image. The band corresponding to the RNase III.RNA complex was gel-excised, fragmented, and digested with 100 µg of proteinase K at 55°C for 2 h (Supplementary Figure 1). RNA was extracted using phenol:chloroform:isoamyl alcohol (PCI) extraction, reverse transcribed using Superscript IV (Thermofisher), and PCR amplified. PCR products were separated on a 2% MetaPhor^TM^ agarose gel (Lonza) and smeared amplicons were gel-excised using the MinElute gel extraction Kit (Qiagen). Libraries were quantified with the Qubit^TM^ dsDNA HS assay kit (Thermofisher), pooled and submitted for HiSeq paired-end 150 bp sequencing at NovogeneAIT Genomics Singapore.

### Annotation of *S. aureus* JKD6008 transcriptome

The ANNOgesic pipeline, which integrated transcriptomics data from RNA-seq, dRNA-seq, and Term-seq, was used to generate a custom annotation of the *S. aureus* JKD6009 genome in a previous report (McKellar *et al*., 2022; Mediati *et al*., 2022). The features in this customized JKD6009 genome annotation were transferred to the *S. aureus* JKD6008 genome using the Rapid Annotation Transfer Tool (RATT) (Otto *et al*, 2011). This generated an embl output file containing the updated S. *aureus* JKD6008 genome annotation which was converted into a GFF format using EMBOSS seqret and custom scripts.

### Analysis of CLASH hybrids

Analysis of the CLASH hybrids was performed similarly as described in our previous report (Mediati *et al*., 2022). Briefly, BBDuk was used to trim the 5’ adapter sequence of the raw data using the parameters ‘ktrim=l k=23 mink=11 hdist=1’ followed by quality trimming with the parameters ‘qtrim=rl trimq=30’(Bushnell, 2014). After trimming, BBmerge was used to merge the paired-end reads using the default parameters(Bushnell *et al*, 2017). Using the pyCRAC pipeline, demultiplexing and removal of PCR duplicates was performed using the pyBarcodeFilter.py and pyFastqDuplicateRemover.py scripts, respectively. Reads were mapped to the *S. aureus* JKD6008 genome using Novoalign (version 2.07) and read counts for each RNA species was calculated using the pyReadCounters.py script. Detection and annotation of the RNA-RNA interactions was performed using the Hyb bioinformatics pipeline and statistical analysis of these RNA-RNA interactions were calculated using custom R scripts that are outlined a previous report (Waters *et al*., 2017).

### Differential gene expression (RNA-seq)

Total RNA from triplicate samples of JKD6008 grown to an OD_600nm_ of 0.8 that were either untreated or treated with 8 µg/mL vancomycin for 30 minutes was harvested using GTC-phenol extraction. RQ1 RNase-free DNase (Promega) was used to deplete genomics DNA and RNA was further purified using PCI extraction. Total RNA samples were sent to NovogeneAIT Genomics Singapore for paired-end 150 bp sequencing on the NovaSeq platform.

Sequencing data was pre-processed using BBtools and mapped to the *S. aureus* strain JKD6008 genome with BBmap as outlined above. Read counts for each feature was computed using bedtools coverage and differential gene expression analysis was performed using DESeq2 (Love *et al*, 2014).

### Ribosome profiling (Ribo-seq)

The ribosome profiling protocol conducted was based off previous studies with some modifications (Choe *et al*, 2019; Latif *et al*, 2015; Oh *et al*, 2011). *S. aureus* cultures were grown to an OD_600nm_ of 0.8 in MH media and then treated with 100 µg/mL chloramphenicol for 2 min, harvested and snap frozen in liquid nitrogen. Cell pellets were resuspended with 1 mL of cold lysis buffer (10 mM CaCl_2_, 100 mM NH_4_Cl, 20 mM Tris-HCl pH 8.0, 0.1 % NP40, 0.4% Triton X-100, 10 U/ml RQ1 DNase (Promega), 1 mM Chloramphenicol, 100 µg/mL lysostaphin) and lysed using 3 V of zirconia beads in a FastPrep-24^TM^ 5G homogenizer (MP Biomedicals) for two cycles of 40 seconds at 6.5 m/sec with 1-minute rest in between. Samples were clarified by centrifugation and 200 µL of supernatant was treated with 1000 U of micrococcal nuclease (MNase) (Sigma-Aldrich) supplemented with a final concentration of 10 µM CaCl_2_ and incubated at 25°C shaking at 250 rpm for 1 h. Polysomes were purified using Illustra Microspin S-400 HR columns (Cytiva).

Ten micrograms of RNA were resolved on a 12% TBE-Urea polyacrylamide at 200 V for at least 4 hours in 1x TBE buffer and 20-50 nt bands were gel-excised, fragmented, and recovered. The 3′ ends of purified RNA were dephosphorylated with the addition of T4 PNK enzyme (New England Biolabs) and 10 mM ATP was added to phosphorylate the 5’ ends of the RPF (ribosome protected fragments). RPF were purified using PCI extraction and unique barcoded 5’ linkers were ligated to the purified RPF. The 5’ end ligated samples were purified using PCI extraction and then the 3’ App-PE adapters were ligated to the RPF using 40 U T4 RNA ligase I (NEB). The 5’ and 3’ ligated samples were resolved on a 12% TBE-Urea polyacrylamide gel and the band corresponding to the RPF was gel-excised, fragmented, and recovered. Purified RPFs were reverse transcribed using Superscript III (Thermofisher) and rRNA depletion was performed using the Zymo-Seq RiboFree® Total RNA Library Kit (Zymo Research) with two modifications: the first-strand cDNA synthesis and adapter ligation steps were omitted. The cDNA was rRNA depleted at 68°C for 14 h, PCR amplified, separated on a 2% MetaPhor^TM^ agarose gel (Lonza) and then gel-excised using the MinElute gel extraction Kit (Qiagen). Libraries were quantified with the Qubit^TM^ dsDNA HS assay kit (Thermofisher), pooled and submitted for HiSeq paired-end 150 bp sequencing at NovogeneAIT Genomics Singapore.

### Protein extraction and mass spectrometry (LC-MS/MS)

*S. aureus* cultures were grown to an OD_600nm_ of 0.8 in MH media and treated with 8 µg/mL vancomycin for 30 minutes. Cultures were harvested by centrifugation and cells pellets were resuspended in 1 mL of lysis buffer (50 mM Tris–HCl (pH 7.8) (Sigma-Aldrich), 150 mM NaCl, and 1 tablet “cOmplete” EDTA-free protease inhibitor (Roche) per 50 mL). The cell resuspension was lysed with 3 V of zirconia beads in a FastPrep-24^TM^ 5G homogenizer (MP Biomedicals) for two cycles of 40 seconds at 6.5 m/sec with 1-minute rest in between. Cell lysates were cleared by centrifugation and protein concentration was quantified using the Qubit™ Protein and Protein Broad Range Assay Kit (ThermoFisher). Protein samples were sent to the UNSW Bioanalytical Mass Spectrometry Facility for liquid chromatography-mass spectrometry (LC-MS/MS) processing and analysis using the Orbitrap Fusion Lumos Tribrid Mass Spectrometer (ThermoFisher).

Raw mass spectrometry data was processed using the MaxQuant program(Cox & Mann, 2008). The Mascot search engine was used to match peptides in the raw mass spectrometry data against the whole *S. aureus* reference proteome set in the UniProt database(Consortium, 2007). Using Perseus, contaminants were filtered and label-free quantification (LFQ) values were log_2_ transformed. Peptides that did not contain LFQ-intensity values in at least 70% of total samples were removed and missing values were imputed using a width and downshift values of 0.3 and 1.8, respectively. The difference in log_2_(LFQ) values between the control and vancomycin-treated group was then calculated(Tyanova *et al*, 2016). The statistical significance of these log_2_(LFQ) differences was calculated using a two-tailed student t-test and then FDR corrected using the Permutation-based FDR method.

### Data clustering (SOMS)

*Data normalization and compilation of multi-omics data table*. The untreated samples were analyzed using RNA-Seq, Ribo-Seq and Label Free Quantification (LFQ) proteomics. The transcriptomics and proteomics datasets both had three biological replicate samples each for the untreated controls, while the Ribo-seq experiment had biological duplicate samples. Each biological replicate sample were log_2_ normalized, and the average value from the biological replicate samples and from each type of -omics dataset were calculated independently. The resulting averaged values were further normalized using the z-normalization to calculate the standard score. The standard score represents the number of sample standard deviation by which the average log-transformed abundance value of a single transcript or protein is above or below the mean value for the sample. A kernel density plot and normal Q-Q plot was generated to ensure that the data appears normally distributed after z-score normalisation. The standard scores were combined into a single multi-omics data table, with transcript, ribosome occupancy, and the corresponding protein product as rows and the samples from different types of -omics and treatments as columns.

*Enrichment of binding targets of sRNA from co-regulated clusters of transcripts and protein products.* Self-organizing maps (SOMs) were used to identify co-regulated clusters of transcripts and protein products from the multi-omics data. Here, the SOMs function from the R package Kohonen v 3.0.11(Wehrens & Buydens, 2007) was used to divide *S. aureus* genes into three clusters based off their gene expression profile across all untreated samples using the single multi-omics data table and JKD6008 genome annotation GFF file as inputs. The co-regulated transcript and/or protein products from each cluster were analyzed for enrichment of verified sRNA-mRNA interactions in *S. aureus* and RNase-III CLASH interactions using Fisher’s exact test. For each cluster which contained RNase III-CLASH interactions, a metagene plot of the cumulative count of RNA-RNA interactions relative to the start codon was generated using a custom script.

### Construction of GFP-translational fusions

Three target mRNAs *lip*, *SAA6008_01130*, and *hpr* were PCR amplified from the JKD6008 gDNA using Phusion Hot Start Flex Polymerase (New England Biolabs) with primers that incorporate the BgIII and EcoRV restriction sites at the 5’ and 3’ ends, respectively. PCR amplicons were cloned into pCN33p::ptufA-isaA-GFP by switching out the *isaA* insert between the BglII/EcoRV sites. Small RNAs of interest (RsaOI, RNAIII, and RsaE) were amplified with primers that incorporate the PstI and EcoRI restriction sites at the 5’ and 3’ ends, respectively. Purified amplicons were cloned into a pICS3::P_tufa_ vector (Ivain *et al*, 2017) and after verifying the constructs by Sanger sequencing, the pICS3::P_tufa_ construct was transformed into electrocompetent *S. aureus* RN4220 and overnight cultures were stored at - 80°C with 16% (v/v) glycerol. To generate co-transformants, the transformed strains were then made electrocompetent and subsequently transformed with the pCN33 vector.

### Flow cytometry measurement of GFP expression

*S. aureus* RN4220 strains co-transformed with the target GFP-translational fusion and sRNA expression plasmid were streaked onto BHI agar supplemented with 10 µg/mL chloramphenicol and erythromycin. For each sample, three biological replicates were individually inoculated into 1 mL of 0.22 µM filtered BHI media and incubated overnight at 37°C with 200 rpm shaking. Cultures were subsequently normalised to an OD_578nm_ of 2 using 0.22 µM filtered BHI media and fluorescence was quantified with the LSRFortessa Special Order Research Product cell analyser (BD Biosciences) using a 530/30 nm bandpass filter. The forward scatter, side scatter, and background cellular fluorescence were gated initially on a wild-type RN4220 population and at least 100,000 events were sampled. The median fluorescence intensity (MFI) for each biological replicate was determined using the FlowJo software (version 8) and a standard two-tailed student’s t-test was performed to determine the difference in mean MFI between samples.

### Antibiotic sensitivity testing by spot dilution

*S. aureus* strains JKD6009, JKD6008, and Mu50 containing the pSD1 constructs were inoculated into 5 mL of MH media supplemented with 10 µg/mL chloramphenicol and incubated overnight at 37°C with 200 rpm shaking. Cultures were then normalised to an OD_600nm_ of 1, aliquoted into a 96-well microtitre plate (Corning^®^), serially diluted (up to 10^-7^) in MH media and spotted onto solid MH plates supplemented with 500 ng/mL aTC, 5 µg/mL chloramphenicol, and with 0, 1, 2, or 3 μg/mL vancomycin. Spot plates were air dried at room temperature and then incubated at 37°C for 24-48 h. Results were imaged on the Bio-Rad Chemi-doc using the default trans-white light settings.

### Antibiotic sensitivity testing by growth curve analysis

Growth curve analysis of *S. aureus* strains JKD6008, JKD6009 and other strain derivatives in the presence of different antibiotics were measured using the Bioscreen C MBR (Growth Curves USA, Piscataway, NJ). Here, single colonies were inoculated into 5 mL of MH media and incubated at 37°C shaking at 200 rpm overnight. Overnight cultures were normalised to an OD_600nm_ of 2 and biological triplicates were diluted 1/100 into three wells containing 200 µL of MH media supplemented with either 0, 2, 3, or 4 µg/mL of vancomycin. Three wells containing only MH media were used as blanks. The plate was incubated in the Bioscreen C at 37°C with continuous low shaking and OD_600nm_ readings were recorded every 20 minutes for 24 hours. Growth curve data was analysed using GraphPad Prism.

### Northern blot analysis

Total RNA purified (2 µg) from GTC-phenol: chloroform extraction was resolved on a 12% TBE-Urea polyacrylamide gel and electrophoresed at 200 V for at least 4 hours in 1x TBE buffer. The gel was post-stained with SYBR Safe DNA Gel Stain and intact RNA was confirmed using the Bio-Rad Chemi-doc. The RNA was transferred onto a nylon membrane (GE Healthcare Life Sciences) soaked in 0.5X TBE buffer via electrophoresis at 30 V for 4 hours at 4°C and then crosslinked in a Strategene Auto-crosslinker with 1200mJ UV-C. The membrane was pre-hybridised in ULTRAhyb-Oligo buffer (Thermofisher) at 37°C for 30 minutes. The 35 mer DNA oligonucleotide probes were radiolabelled with ^32^P-ATP at 37°C for 1 hour and then purified over a G50 size exclusion column (GE Healthcare Life Sciences) following the manufacturer’s instructions. The purified oligonucleotide mix was added to the nylon membrane and hybridised at 37°C overnight. The membrane was washed four times in wash buffer (2X SSPE buffer (Sigma-Aldrich) and 0.1% SDS) at 37°C for 30 minutes. Afterwards, the membrane was exposed on a BAS Storage Phosphor Screen and detection of the hybridised probe was performed on the LAS-3000 (Fujifilm).

## RESULTS

### RNase III-CLASH uncovers the VISA sRNA interactome

In our earlier analysis we used RNase III-CLASH to profile RNA-RNA interactions associated with RNase III in the methicillin-resistant *S. aureus* isolate JKD6009 (McKellar et al., 2022; Mediati et al., 2022). In an effort to capture sRNA interactions that contribute to vancomycin tolerance we have applied RNase III-CLASH to the clinical VISA strain JKD6008. To facilitate the purification of UV-crosslinked RNA-protein complexes from VISA, we first constructed a chromosomal translational fusion of RNase III with a C-terminal linked His_6_-TEV-FLAG dual affinity tag in *S. aureus* VISA strain JKD6008. JKD6008 has a thicker cell wall compared to the JKD6009 parent strain (Howden *et al*., 2010), and our initial RNase III purifications were unsuccessful. We found that the inclusion of 10 µg/mL lysostaphin and lysis for two cycles of 40 seconds using a FastPrep-24TM 5G homogenizer (MP Biomedicals) increased cell lysis and recovery of RNase III in the soluble fraction, but the harsher lysis conditions also resulted in the release of an approximately 50 kDa protein that bound to the M2 FLAG resin. LC-MS/MS analysis indicated that the FLAG purified protein was the immunoglobulin-binding protein, Sbi. To remove Sbi and improve RNase III recovery, we incubated our cell lysates with mouse IgG agarose to sequester Sbi. After removing Sbi from the cell lysate, we were able to purify RNase III-HTF complexes from VISA cell lysates.

Biological duplicates of RNase III-CLASH libraries were prepared from *S. aureus* JKD6008 and the isogenic His-TEV-FLAG tagged strain *rnc*-HTF as described previously (excepting improved lysis conditions described above) (Mediati *et al*., 2022). Briefly, cultures grown to an OD_600nm_ of 0.8 were treated with and without 8µg/ml vancomycin for 10 minutes. Cultures were then immediately crosslinked with 400 mJ of UV-C (Vari-X-Link, UV03), harvested by vacuum filtration, and lysed as described above. RNase III-associated RNAs were affinity purified under denaturing conditions and linkers ligated to RNA-protein complexes followed by cDNA library construction.

RNase III associated RNAs were sequenced and analysed using the snakemake pipline Hyb-CRAC-R that incorporates the pyCRAC package (Webb *et al*, 2014) for identifying protein binding sites, *hyb* (Travis *et al*, 2014) that identifies RNA-RNA interactions, and our statistical analysis of RNA hybrids in the CLASH dataset. From these analyses, 775 statistically significant RNA-RNA interactions were recovered from VISA strain JKD6008 including 170 sRNA-RNA interactions (Figure 1). To assign RNA fragments recovered by CLASH to genomic features, the genome annotation previously generated for the parental MRSA strain JKD6009 was transferred to the isogenic VISA isolate JKD6008 using RATT (Otto *et al*., 2011). This annotation incorporated RNA boundaries determined by dRNA-seq and Term-seq, and includes experimentally determined mRNA 5’ and 3’ UTRs. The 755 RNA-RNA interactions in VISA covered a broad range of RNA classes (Figure 2A) and consistent with earlier work, mRNA-mRNA interactions are the most recovered interaction type supporting the idea that *S. aureus* may utilise regulatory mRNA interactions to control gene expression (Mediati *et al*., 2022). A metagene analysis showed that both RNA-RNA and sRNA-RNA interactions were enriched at the start codon, which corresponds to the canonical mechanism of sRNA regulation by occluding the RBS (Figure 2B). Comparison of the VISA and VSSA sRNA-RNA interactomes revealed that 11 sRNA-RNA interactions are shared between the MRSA and VISA interaction networks, seven of which were have been experimentally validated including RsaA-RNAIII, RNAIII-*esxA*, RsaOG-RsaE, SprX2-*spoVG*, RsaA-*mgrA*, and RsaA-*ssaA* (Eyraud *et al*, 2014; McKellar *et al*., 2022; Tomasini *et al*, 2017)(Figure 2C). The small amount of overlap between these datasets likely reflects the small size of the VISA dataset and sampling of a large sRNA interactome rather than a large divergence in the VISA and VSSA interaction networks.

**Figure 1.**
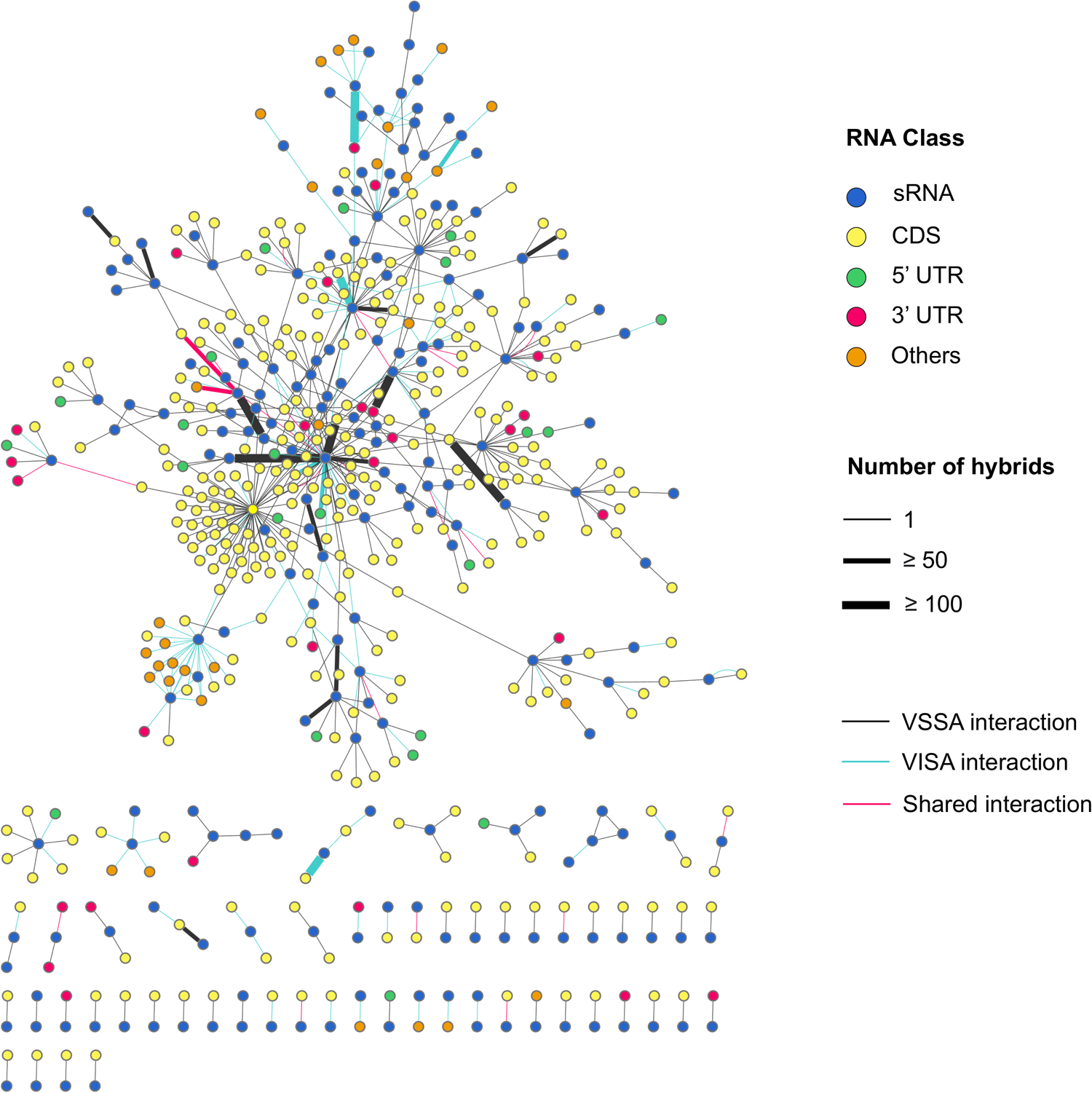
Small RNA interactome of *S. auereus* strains JKD6008 (VISA) and JKD6009 (VSSA). Each RNA is represented as a node (circles) and are coloured according to their RNA class as indicated by the legend (top right). RNA-RNA interactions (FDR<0.05) are represented as edges (lines) and the thickness of each edge represents the hybrid count for each RNA-RNA interaction. Individual RNA-RNA interactions captured in JKD6008 and JKD6009 are represented as blue and black edges, respectively (bottom right).

**Figure 2.**
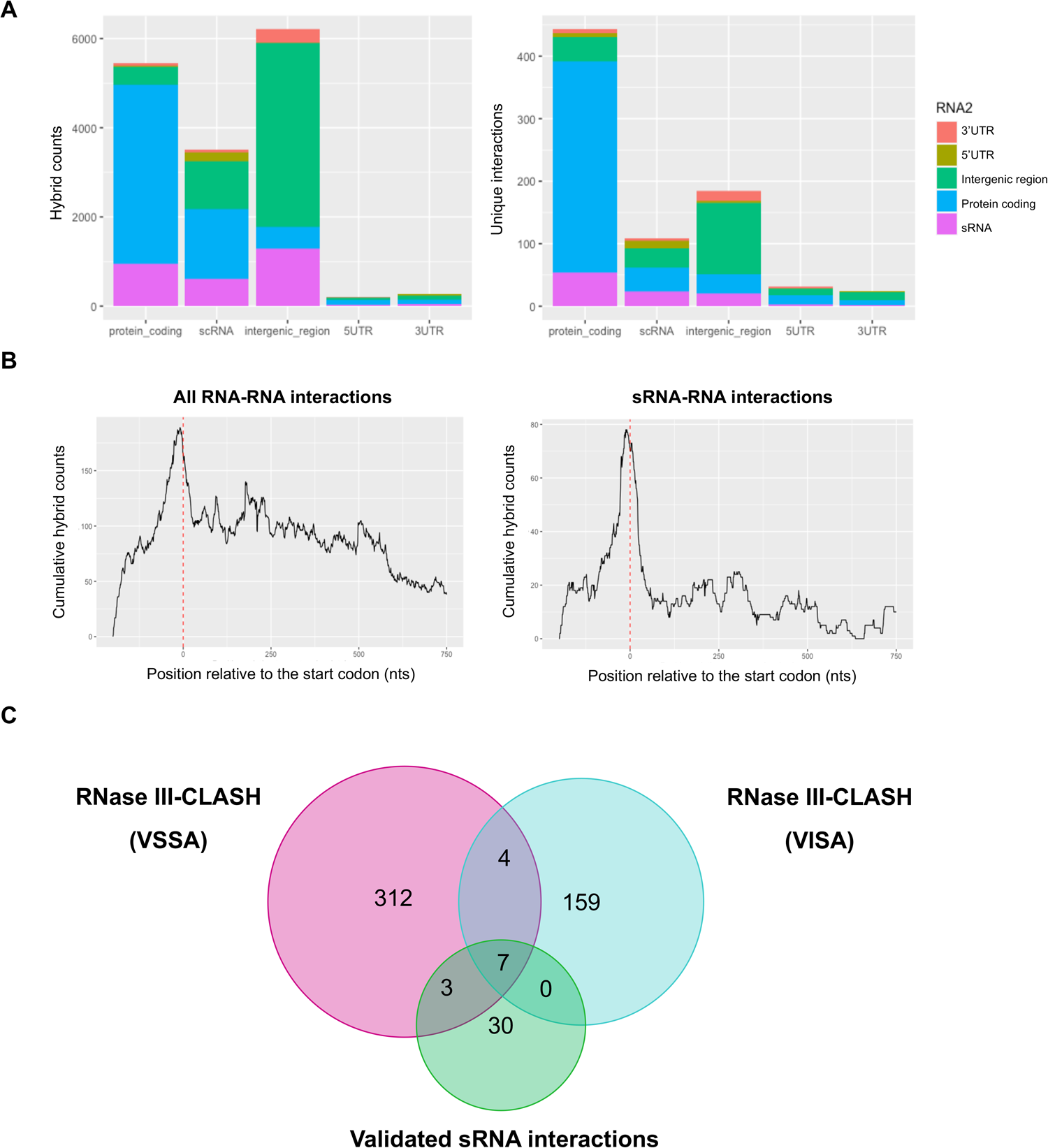
RNA-RNA interactions recovered by RNase III-CLASH in vancomycin tolerant *S. aureus* str. JKD6008 (VISA). (A) Histogram of RNA classes recovered by JKD6008 RNase III-CLASH. The total number of hybrid reads (PCR duplicates collapsed, left panel) and the number of unique RNA-RNA interactions (right panel). RNA classes for hybrid read halves are indicated (right). Note that multiple hybrid reads can represent a single (unique) RNA-RNA interaction. **(B)** Metagene plot of the cumulative count of RNA-RNA (left) and sRNA-RNA (right) interactions relative to the mRNA start codon as indicated by the red dashed line. **(C)** Venn diagram of interactions shared between the sRNA interactomes of VISA (JKD6008), VSSA (JKD6009), and experimentally validated *S. aureus* sRNA-mRNA interactions from the literature.

To determine the potential biological functions of sRNAs, the VISA and MRSA sRNA interactomes were merged and each sRNA node was extracted for GO term analysis of the associated mRNAs. GO term analysis revealed that 26 sRNA clusters were enriched for GO terms associated with several biological processes including purine ribonucleotide synthesis, methylation, and hexose metabolism (*p*<0.05) (Supplementary Table 3).

### Clustering reveals post-transcriptionally regulated mRNAs with RNase III-CLASH sRNA-mRNA interactions

Our RNase III-CLASH data provides a snap-shot of the VISA sRNA interactome but does not inform on the function of the sRNA-mRNA interaction (eg: activation, repression, transcript degradation, translational repression). Further, a proportion of experimentally verified sRNA-mRNA interactions recovered by proximity-dependant ligation are reported to not affect expression of the target mRNA (Faigenbaum-Romm *et al*., 2020; Iosub *et al*., 2020).

We aimed to identify sRNA-mRNA interactions that activate or repress mRNA translation by using a multi-omics approach. Previous work indicates a strong correlation between mRNA and protein abundance within the cell. Many genes that deviate from this correlation are known targets of post-transcriptional regulators (Balakrishnan *et al*, 2022). To identify genes that are being post-transcriptionally regulated, we collected RNA-seq, Ribo-seq, and proteomics data for *S. aureus* strain JKD6008 (VISA) during mid-exponential growth in Muller-Hinton media. For each biological replicate sample from the RNA-seq, Ribo-seq and proteomics datasets the average expression value for each condition was calculated across biological replicates in each ‘omics datatype. We found a high level of correlation between our RNA-seq and Ribo-seq datasets (Pearsons correlation 0.8 from 2679 genes) and lower correlation between RNA-seq and proteomics data (Persons correlation 0.32 from 566 genes) (Supplementary Figure 2).

To convert the distributions of values to the same scale, the average expression values for each -omics dataset were transformed to z-scores and the datasets were confirmed to be normally distributed (Supplementary Figure 3). Self-organising maps (SOMS) was used to cluster genes with similar z-score normalised transcriptional and translational values (Kohonen, 2013; Tamayo *et al*, 1999). We used to 39 experimentally verified *S. aureus* sRNA-mRNA interactions from past literature (Supplementary Table 4) to determine if we could enrich functional sRNA-mRNA interactions using SOMS clustering (Figure 3). Three clusters of genes with different transcriptional and translational patterns were generated using SOMS. Cluster 1 is enriched for verified translationally repressive sRNA-mRNA interactions (9/35 within cluster 1, Fishers exact test *p* =0.0198)(Supplementary Figure 4A). The 173 genes in cluster 1 exhibit a pattern of average to high transcript abundance, but lower protein levels, suggesting that the genes in this cluster are being post-transcriptionally repressed. The 187 genes in cluster 2 have transcript abundance that corresponds with protein expression, while the 207 genes in cluster 3 appear to collectively have moderately higher protein levels relative to transcript abundance (Figure 3).

**Figure 3.**
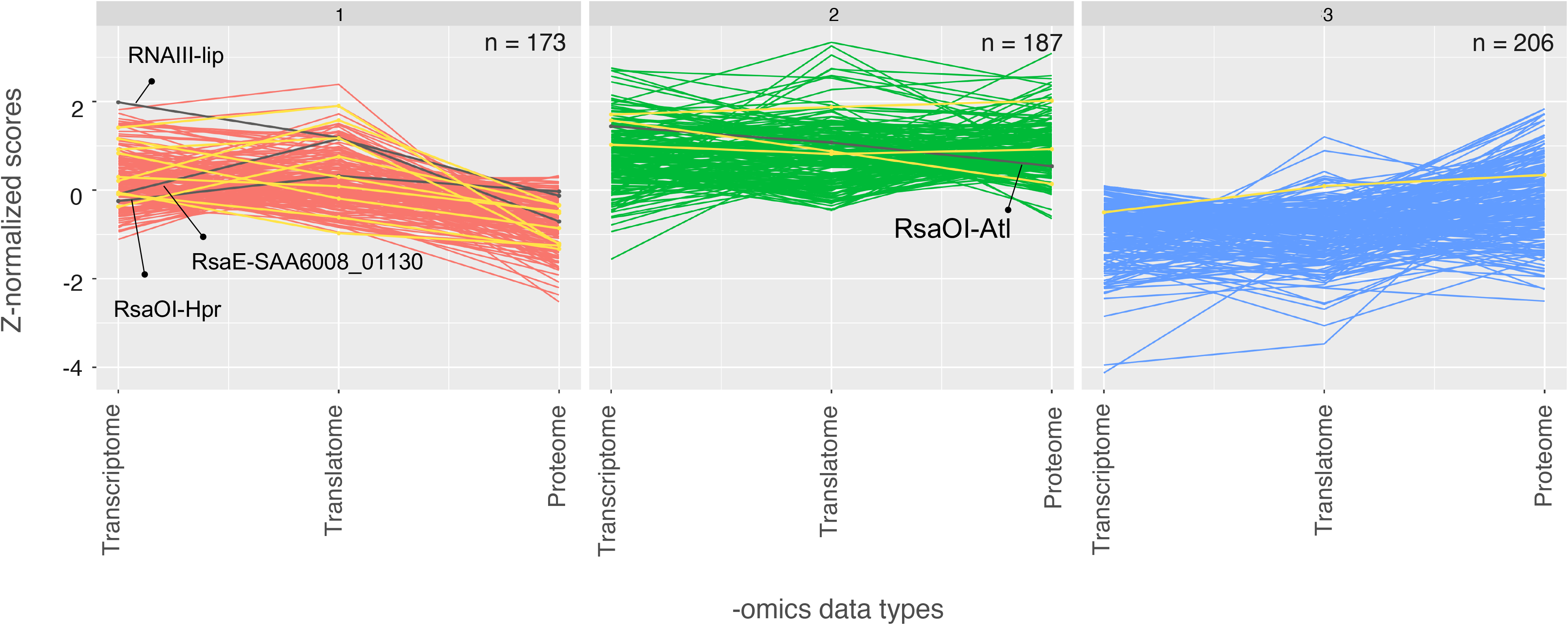
Self-organising maps (SOMS) reveals a cluster of genes enriched with translationally repressive sRNA-mRNA interactions. Each graph displays a SOMS cluster of genes with similar transcriptional and translational patterns. The x-axis contains the - omics data types for RNA-seq (transcriptome), Ribo-seq (translatome), and mass spectrometry (proteome) under the control conditions. The y-axis contains the z-normalised scores for all the -omics data types. The number of genes present in each cluster (from a total of 566 genes with signal in all datasets) is indicated (n) in the top right of each plot. Verified *S. aureus* sRNA-mRNA interactions are indicated in yellow and four RNase III-CLASH interactions tested in this study are indicated in black.

Next, we cross-referenced the 412 *S. aureus* sRNA-mRNA interactions captured from both the VISA and VSSA sRNA interactome with the three SOMs clusters (Figure 3, Supplementary Figure 4B). Both cluster 1 and 2 are enriched for sRNA-mRNA interactions captured in our RNase-III CLASH network (Fishers exact test, *p* < 0.0001).

As cluster 1 has lower than expected protein abundance and verified repressive sRNA-mRNA interactions, we speculated that this cluster is enriched for translationally repressive RNase III-CLASH interactions. We examined whether cluster 1 interactions are enriched around the 5’ UTR and start codon since sRNAs canonically interact with this region to repress translation. Metagene plots of the subset of RNase-III CLASH sRNA-mRNA interactions detected in each cluster were used to determine which regions of the mRNA were being targeted for sRNA regulation. For the 88 sRNA-mRNA interactions in cluster 1, the mRNA read halves predominantly mapped to the 5’ UTR and start codon (Supplementary Figure 4Ci). A slightly less pronounced enrichment was recovered for clusters 2 and 3 (Supplementary Figure 4Cii&iii).

Previous work indicated that functional sRNA-mRNA interactions are more likely to contain a higher number of hybrid reads (Faigenbaum-Romm *et al*., 2020). We examined the number of hybrid reads recovered for sRNA-mRNA interactions in each cluster (Supplementary Figure 5). While cluster 1 contains many of the highest abundance interactions, no statistically significant difference was observed between the three clusters.

Collectively our data indicates that clustering of genes with lower than expected protein abundances enriches for experimentally verified and CLASH-recovered sRNA-mRNA interactions.

### SOMs cluster 1 contains sRNA-mediated translationally repressed genes

Our SOMs analysis suggests that sRNA interactions with mRNAs in cluster 1 may be responsible for translational repression of the gene (lower than expected protein abundance). To evaluate their function, three sRNA-mRNA interactions from our RNase-III CLASH network that were present in cluster 1 were tested using GFP-translational fusions. These three interactions include RNAIII-*lip*, RsaE-*SAA6008_01130*, and RsaOI-*hpr*. Each target mRNA was cloned into the GFP-translational fusion vector pCN33::P*_tufA_*. Each mRNA was cloned from the +1 transcriptional start site identified by dRNA-seq and included the native Shine-Dalgarno sequence that overlaps the predicted CLASH interaction site. These were transformed into *S. aureus* RN4220 strains containing the pICS3::P*_tufA_*vector constitutively expressing their cognate regulatory sRNA and fluorescence was measured using a flow cytometer. Transcription of the sRNAs RNAIII and RsaE reduced the fluorescence signal of the triacylglycerol lipase precursor *lip* and *SAA6008_01130*, which encodes an acetyltransferase family protein, by 2.05 and 3.10-fold (both *p*< 0.05), respectively (Figure 4A and B). RsaOI reduced *hpr* expresson by 5.16-fold (*p*<0.05) (Figure 4C). All three sRNA-mRNA interactions significantly reduced target mRNA translation.

**Figure 4.**
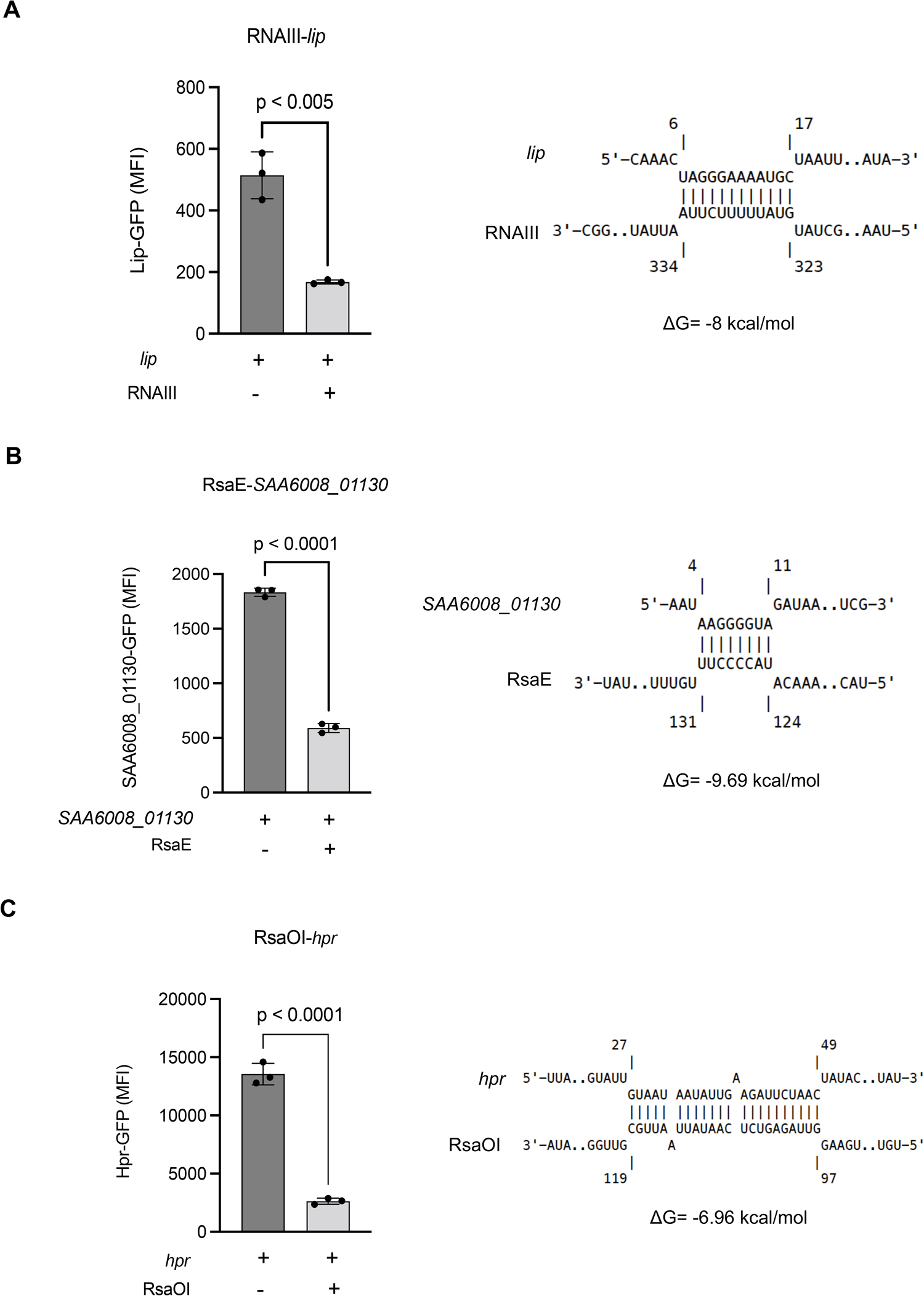
Translationally repressive RNase III-CLASH sRNA-mRNA interactions in SOMS cluster 1. GFP-translational fusions for *lip* (A), SAA6008_01130 (B), and *hpr* (C) were constitutively expressed in *S. aureus* strain RN4220 with or without transcription of their cognate sRNAs (indicated above). IntaRNA predicted interactions at mRNA site recovered by RNase III-CLASH and the associated free energy change are indicated on the right. Median fluorescence intensity was measured using flow cytometry and error bars indicate standard error. P values were calculated using a two-tailed *t*-test.

Collectively our data suggest that translationally repressive sRNA-mRNA interactions are enriched in clusters with a lower than expected protein abundance. The correlation in transcript and protein abundance may provide a useful tool to identify post-transcriptionally regulated genes and functional sRNA-mRNA interactions.

### The small RNA RsaOI is induced by vancomycin

We next looked to use our VISA CLASH network and meta-omics clustering to identify sRNAs and interactions that contribute to vancomycin tolerance. We performed RNA-seq on VISA samples that were treated with 8 µg/mL vancomycin for 30 minutes (Figure 5A). RsaOI was the highest upregulated sRNA in response to vancomycin treatment (log_2_FC= 5.91, aka Teg47 or Sau6477). Differential RNA sequencing (dRNA-seq) and Term-seq data, were used previously to map the transcriptional start sites and termination sites in *S. aureus* JKD6009 genome (Mediati et al. 2022) and indicate that RsaOI is 292 nt long transcript (Figure 5B), which is slightly larger than earlier northern analysis (Bohn *et al*, 2010). Northern blot analysis demonstrated a dose-dependent induction of RsaOI in response to vancomycin treatment (Figure 5C). The vancomycin-responsive WalKR two-component system is an important contributor to the VISA phenotype. To determine whether RsaOI expression is under the control of WalKR, northern blot analysis for RsaOI was performed in wild-type USA300 strain NRS384, Δ*yycHI*, and *walK*^H271Y^ mutants with and without 8 µg/mL vancomycin. RsaOI was induced similar to wild-type in the *walKR* mutants, indicating that it is not regulated by WalKR (Supplementary Figure 6). To test vancomycin tolerance, JKD6008 and Δ*rsaOI* were grown in MH media in the presence or absence of sub-MIC vancomycin (3 μg/mL). These results indicate that *rsaOI* is strongly induced by vancomycin but is not required for vancomycin tolerance (Supplementary Figure 7).

**Figure 5.**
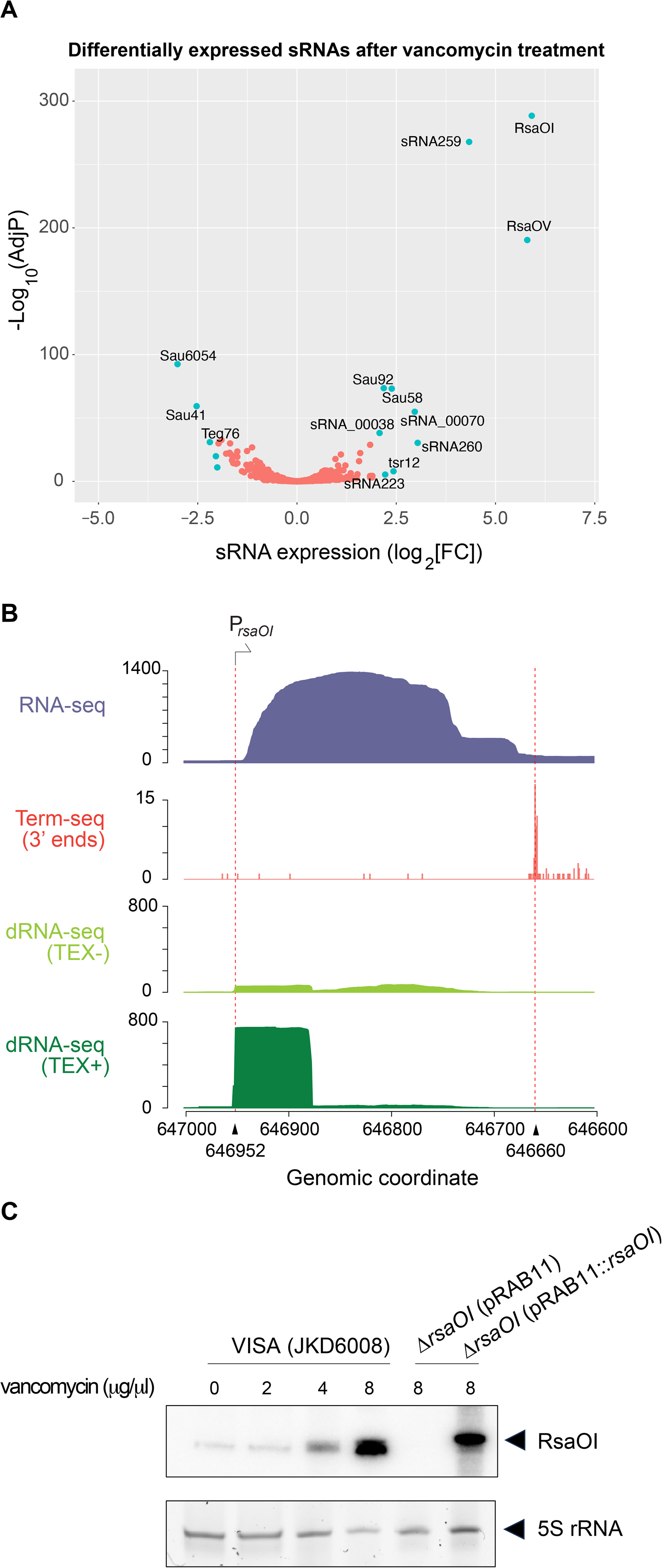
RsaOI is induced by vancomycin stress. **(A)** Volcano plot of differentially expressed sRNAs in JKD6008 after 30 minutes treatment with 8μg/μL vancomycin treatment. Small RNAs with log_2_FC >=|2| and FDR<0.05 are indicated in light blue. **(B)** 5’ and 3’ end mapping of RsaOI. From top to bottom, total RNA-seq reads (blue), Term-seq RNA 3’ end reads after vancomycin treatement (red), dRNA-seq reads after vancomycin treatment (no TEX treatment), dRNA-seq reads after vancomycin treatment (with TEX treatment). Dashed red lines indicated RsaOI 5’ and 3’ ends determined by dRNA-seq and Term-seq. **(C)** Northern blot analysis of RsaOI expression after 30 minutest treatment with increasing vancomycin concentrations (indicted above) in VISA (JKD6008) and the isogenic *ΔrsaOI* mutant. Migration of RsaOI is indicated by the black arrow. Sybr stained 5S rRNA is included as a loading control.

### RsaOI controls the expression of the major autolysis Atl critical to vancomycin tolerance

To identify functional targets of RsaOI we looked within our CLASH data using the SOMS clustered mRNA targets. RsaOI has 9 differentially expressed RNase III-CLASH mRNA targets (Figure 6A) and five of these are downregulated during treatment with vancomycin using a log_2_FC cut-off >= |0.5| (Figure 6B), suggesting that the RsaOI upregulation causes the repression of these mRNA targets. To identify functional RsaOI targets we repeated our SOMS clustering on RNA-seq and proteomics data and included vancomycin treated VISA cultures (Figure 6C). We segregated the data into three clusters that separated the ‘omics data into distinct profiles. Cluster 3 is enriched for experimentally verified sRNA targets and has lower protein abundance than expected from mRNA levels. We overlaid our RNase III-CLASH network. One RsaOI mRNA target (*hpr*) falls within cluster 3 and we had previously confirmed that this interaction is functional (above). Interestingly, three targets fall within cluster 1 which shows correlated RNA and protein levels. Within this cluster, SAA6008_01438 displays higher protein abundance compared to transcript levels. We focused on the mRNA *atl* in cluster 1 that encodes a bifunctional autolysin involved in cell wall turnover and cell division as this has previously be implicated in glycopeptide tolerance (Nega *et al*, 2020; Zheng *et al*, 2022). RNase III-CLASH predicted that RsaOI binds to the coding region of *atl* mRNA consistent with mRNA decay rather than translational repression as the mechanism of action (Figure 6C). After treatment with vancomycin, *atl* mRNA and protein are downregulated to a similar extend consistent with the profile of cluster 2 (Figure 6B and D). We used a GFP-translational fusion to evaluate the function of RsaOI on *atl* gene expression. The transcription of RsaOI reduced the fluorescence signal of Atl-GFP by 1.63-fold (*p< 0.05) confirming that the interaction is functional. The RNase III-CLASH interaction indicates that RsaOI base pairs 226 nt downstream of the start codon of *atl* mRNA (Figure 6E). We speculate that RsaOI recruits RNase III to the coding sequence of *atl* mRNA and drives repression through mRNA degradation resulting in more correlated mRNA and protein abundance.

**Figure 6.**
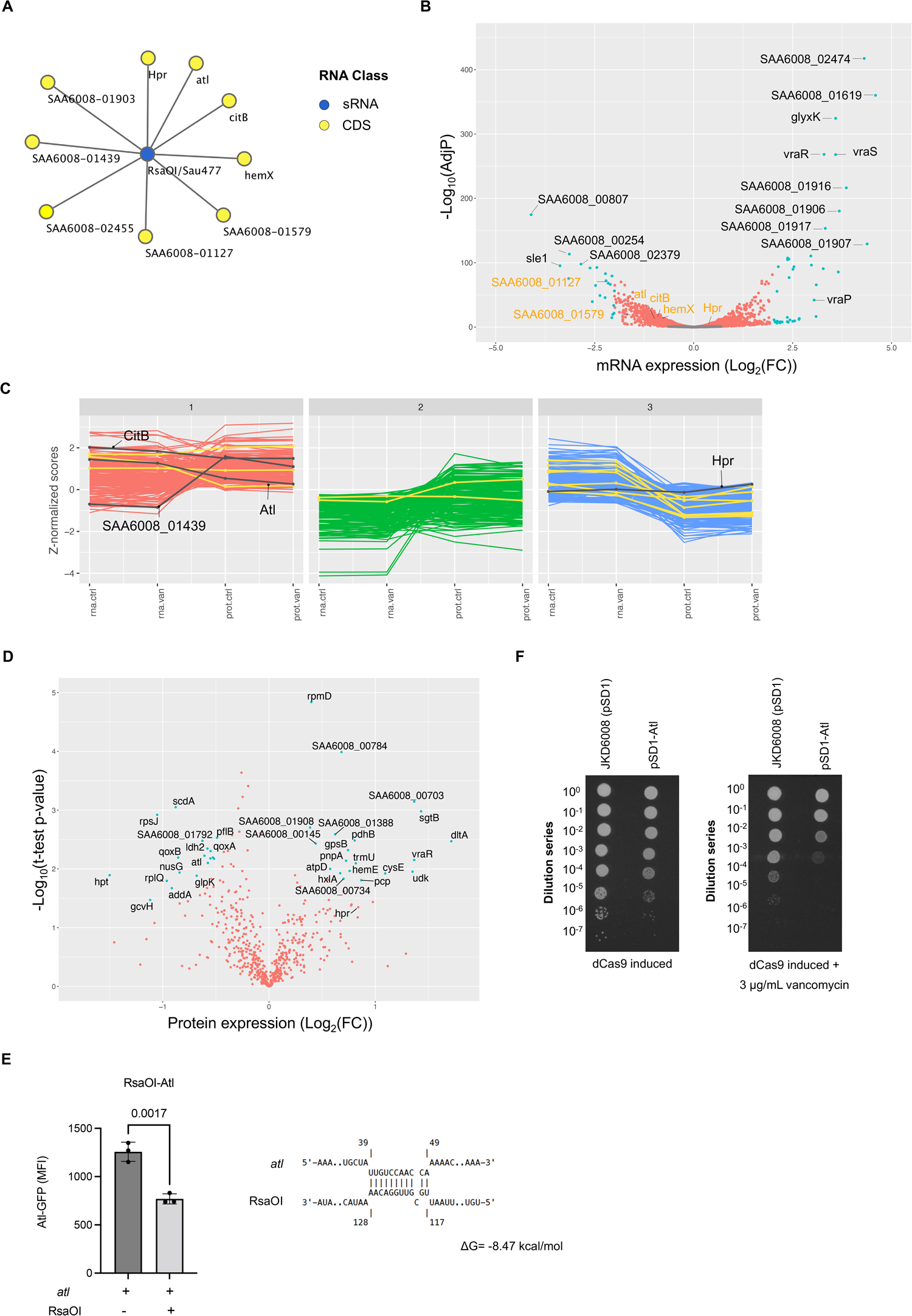
RsaOI mediates the expression of the autoylsin Atl. **(A)** The sub-network of RNase III CLASH RNA-RNA interactions with RsaOI (blue node). Yellow nodes indicate mRNA tagets (indicated). **(B)** Volcano plot of differentially expressed mRNAs in vancomycin treatment (8 µg/mL for 30 minutes) VISA strain JKD6008. RNase-III CLASH identified mRNA targets of RsaOI are highlighted with orange text. Differentially expressed genes that are statistically significant log_2_FC >=|2| and FDR<0.05 are indicated in light blue. **(C)** SOMS clustering of genes with similar transcriptional and translational expression patterns. The x-axis contains the -omics data types for RNA-seq (transcriptome, “rna”) and mass spectrometry (proteome, “prot”) under control conditions (“.ctrl”) or with vancomycin treatment (“.van”). The y-axis contains the z-normalised scores for all the -omics data types. Experimentally validated *S. aureus* sRNA-mRNA interactions are indicated in yellow. RNase III-CLASH identified RsaOI mRNA targets are indicated in black with labels. **(D)** Proteomic analysis of VISA strain JKD6008 with and without vancomycin treatment. Statistical significance was determined using the permutation-based FDR corrected *t*-test and differentially expressed proteins with an FDR <0.05 are indicated in light blue. **(F)** The GFP-translational fusion for the *atl* gene was constitutively expressed in *S. aureus* strain RN4220 with or without transcription of RsaOI (indicated below). RNase III-CLASH predicted an interaction within the coding region of *atl* which is indicated on the right. Median fluorescence intensity was measured using flow cytometry and error bars indicate standard error. P values were calculated using a *t*-test (indicated above plot) **(F)** Spot plate dilution assays to assess vancomycin tolerance in CRISPRi knockdown of *atl* in VISA strain JKD6008.

Atl is downregulated in VISA strains compared to the VSSA parent strains and we assessed vancomycin tolerance of an *atl* knockdown. The *atl* CRISPRi knockdown displayed an approximately 10-fold reduction in growth under the control condition relative to the vector control. In the presence of sub-MIC vancomycin (3 µg/mL), the reduction in growth for the *atl* CRISPRi knockdown increased 100-fold (Figure 6F). This suggests that Atl is required for normal growth and this growth defect is exacerbated in the presence of vancomycin. Increased Atl levels in the *rsaOI* deletion strain may contribute to increased growth under vancomycin stress.

## DISCUSSION

A wealth of sRNA interactome data have been collected in recent years using proximity-dependant ligation techniques including CLASH, RIL-seq, and HiGRIL-seq (Iosub *et al*., 2020; Matera *et al*., 2022; Melamed *et al*., 2016; Waters *et al*., 2017; Zhang *et al*, 2017). While these datasets provide evidence of sRNA-mRNA interactions, they do not indicate the function of the interactions (eg: activating, repressing, target decay, translational control). In addition, it appears that these datasets contain a subset of abundant sRNA-mRNA interactions that do not control target mRNA expression. Discriminating between the different functional sRNA interactions and those with no appreciable effect on gene expression at a systems-level has been challenging (Faigenbaum-Romm *et al*., 2020). Here, we have clustered mRNA targets by their correlation between mRNA abundance, translatome, and proteome in an effort to identify mRNAs that are post-transcriptionally regulated under steady-state conditions. We have applied this approach to identify functional targets of the small RNA RsaOI that is strongly upregulated by vancomycin in the clinical VISA isolate JKD6008.

Earlier work by Faigenbaum-Romm *et al*. (2020) demonstrated that base-pairing strength (ΔG) is a poor predictor of functional sRNA targets from sRNA interactome data, and demonstrated a correlation between Hfq-binding, hybrid read count, and target repression (Faigenbaum-Romm *et al*., 2020). In *Staphylococcus aureus* most sRNAs appear to function independently of the small RNA chaperone Hfq leading to the suggestion that Hfq has been depreciated in *S. aureus* (Bohn et al, 2007) and precluding this approach. We reasoned that post-transcriptional regulation may be reflected in a disconnect between mRNA and protein abundance under steady state conditions. This is supported by recent analysis of the *E. coli* transcriptome and proteome that demonstrates a strong correlation between mRNA and protein abundance (Balakrishnan *et al*., 2022). Indeed, these authors find that most of the genes that do not have a strong correlation are known to be post-transcriptionally regulated, including by small RNAs. Notably, this approach is likely to identify genes controlled by any mechanism of post-transcriptional regulation including through sequence features affecting translation efficiency such as codon usage, RNA secondary structures (eg: RNA thermometers), RNA binding proteins, protein stability, as well as sRNA-mediated regulation (Maier et al, 2009; Nie et al, 2006; Van Assche et al, 2015).

SOMs was used to separate mRNAs into three clusters that have similar transcriptional and translational patterns. Cluster 1 appears to be translationally repressed in that genes have lower than expected protein abundance. Cluster 2 genes have transcript abundance that corresponded with protein expression, and genes in cluster 3 displayed slightly higher protein levels than transcript abundance potentially reflecting translation activation. Using sRNA-mRNA interactions that had previously been described and confirmed, ‘verified interactions’ were enriched in cluster 1 (lower protein levels). We find that CLASH interactions are recovered in all three clusters but are enriched in cluster 1 and cluster 2. Cluster 2 RNA levels correspond to protein levels and for those functional sRNA-mRNA interactions in this cluster we interpret this as protein repression that is primarily driven by mRNA degradation.

Three sRNA-mRNA interactions from cluster 1 were validated using a GFP reporter assay and were found to be strongly repressive. The hybrid read position and predicted base-pairing indicated that these interactions occur around the RBS and start codon region of the target mRNA. To understand where sRNA-mRNA interactions occurred in all clusters, a metagene analysis of the RNase III-CLASH interactions in clusters 1-3 was performed and demonstrated enrichment around the 5’UTR and start codon in clusters 1 and to a lower degree cluster 2 consistent with canonical repression through occlusion of the 30S subunit. However, while cluster 1 contained many of the highest abundance hybrids, no significant difference in mean hybrid counts between the three clusters was found which contrasts with earlier findings by Faigenbaum-Romm *et al*. (2020). These results collectively suggest that cluster 1 contains functional sRNA-mRNA interactions that post-transcriptionally regulate mRNA translation.

Vancomycin intermediate *S. aureus* (VISA) is a major cause of vancomycin treatment failure. Many isolates have a thickened cell wall and reduced autolysis. Single nucleotide polymorphisms (SNPs) that confer the VISA phenotype can be heterogenous and are poorly understood (Cui *et al*, 2003; Howden *et al*., 2010; Muthaiyan *et al*, 2004; Mwangi *et al*, 2007; Pfeltz *et al*, 2000; Renzoni *et al*, 2006; Scherl *et al*, 2006). Previous analysis has suggested that small RNAs may contribute to the VISA phenotype (Howden *et al*., 2013). RsaOI (also termed sRNA131 and Sau6477) was the most upregulated sRNA in response to 8 µg/mL of vancomycin treatment in VISA, consistent with earlier work (Gao *et al*, 2020; Howden *et al*., 2013), and was induced by vancomycin in a dose-dependent manner. RsaOI expression was not regulated by the vancomycin-responsive two-component system WalKR and we hypothesise that RsaOI is controlled by VraSR that is upregulated after vancomycin treatment and controls the divergent *vraABCP* operon (Figure 6B).

To understand the function of RsaOI in *S. aureus*, we examined its mRNA targets. RsaOI has 9 RNase III-CLASH mRNA targets and five of these are downregulated, suggesting that the RsaOI upregulation causes the repression of these five mRNA targets to adapt to antibiotic or cell wall stress. We repeated our SOMS clustering and included RNA-seq and proteomics data from VISA treated with 8μg/ml vancomycin. RsaOI was earlier found to repress the expression of the RNase III-CLASH target *hpr* (phosphocarrier protein Hpr) which was detected in cluster 1 with the majority of experimentally characterised sRNA-mRNA interactions. The RsaOI target *atl* fell within cluster 2 and encodes for a bifunctional autolysin essential for cell wall turnover and cell division (Nega *et al*., 2020; Zheng *et al*., 2022). Atl has been previously implicated in glycopeptide intermediate resistance as it was found to be downregulated in several VISA strains relative to their respective VSSA parent strains. It was postulated that the downregulation of *atl* reduces autolytic activity and contributes to the thickened cell wall phenotype of VISA (Cafiso *et al*, 2012; Howden *et al*., 2011; Wootton *et al*, 2005). In this study, we found that *atl* is downregulated in response to vancomycin in both the RNA-seq and proteomics data and it was present in cluster 1 of our vancomycin-treated SOMs analysis which display a pattern of corresponding transcript and protein levels. We demonstrated that RsaOI expression led to a 1.6-fold reduction in fluorescence signal in the Atl-GFP translation fusion and RNase III-CLASH predicted that RsaOI binds to the coding region of *atl* mRNA. This suggests that RsaOI forms a duplex with the *atl* mRNA that recruits RNase III for *atl* degradation and is consistent with mRNA decay driving reduced protein abundance (correlated mRNA/protein levels). As *atl* was found to be downregulated in several VISA isolates relative to VSSA parent strain (Wootton *et al*., 2005), we reasoned that the knockdown of *atl* would result in increased growth in the presence of vancomycin. However, the *atl* CRISPRi knockdown had reduced growth without antibiotic and the growth defect was exacerbated by the addition of sub-MIC vancomycin. Atl functions in *S. aureus* cell division where it forms a ring at the septum to facilitate partitioning of daughter cells (Nega *et al*., 2020; Yamada *et al*, 1996). It was previously reported that deleting *atl* causes a defect in daughter cell separation (Bose *et al*, 2012) and may explain the mutant phenotype.

Our data collectively suggests that RsaOI is a sRNA that is induced by vancomycin in VISA but is not required for vancomycin tolerance. A similar result was observed by Gao et al. 2020 where deletion of several linezolid induced differentially expressed sRNAs, including RsaOI, yielded no effect on the phenotype (Gao *et al*., 2020). We speculate that RsaOI may be involved in a general cellular stress response (eg: cell wall damage) that is not sufficient to resist vancomycin treatment and/or is functionally redundant.

The approach described here enables the identification of post-transcriptionally regulated mRNAs and coupled with sRNA interactome data provides an initial step towards systems-level functional characterisation of the sRNA-mRNA interaction network. Using these data we have identified several targets of the vancomycin-responsive sRNA, RsaOI and demonstrated that one target, the cell wall autolysin *atl*, contributes to vancomycin tolerance. Our study suggests that the correlation between RNA and protein abundances at steady-state can be used to indicate the function of sRNA-mRNA pairs in sRNA interactome data.

## ACKNOWLEDGEMENTS

The authors thank Tim Stinear and Ian Monk for providing *S. aureus walKR* mutant strains. WW was supported by an Australian Research Training Program scholarship. JJT, DGM, and WW are supported by grants from the National Health and Medical Research Council (NHMRC GNT1139313) and the Australian Research Council (ARC DP220101938).

## SUPPLEMENTARY FIGURE LEGENDS

**Supplementary Figure 1.**
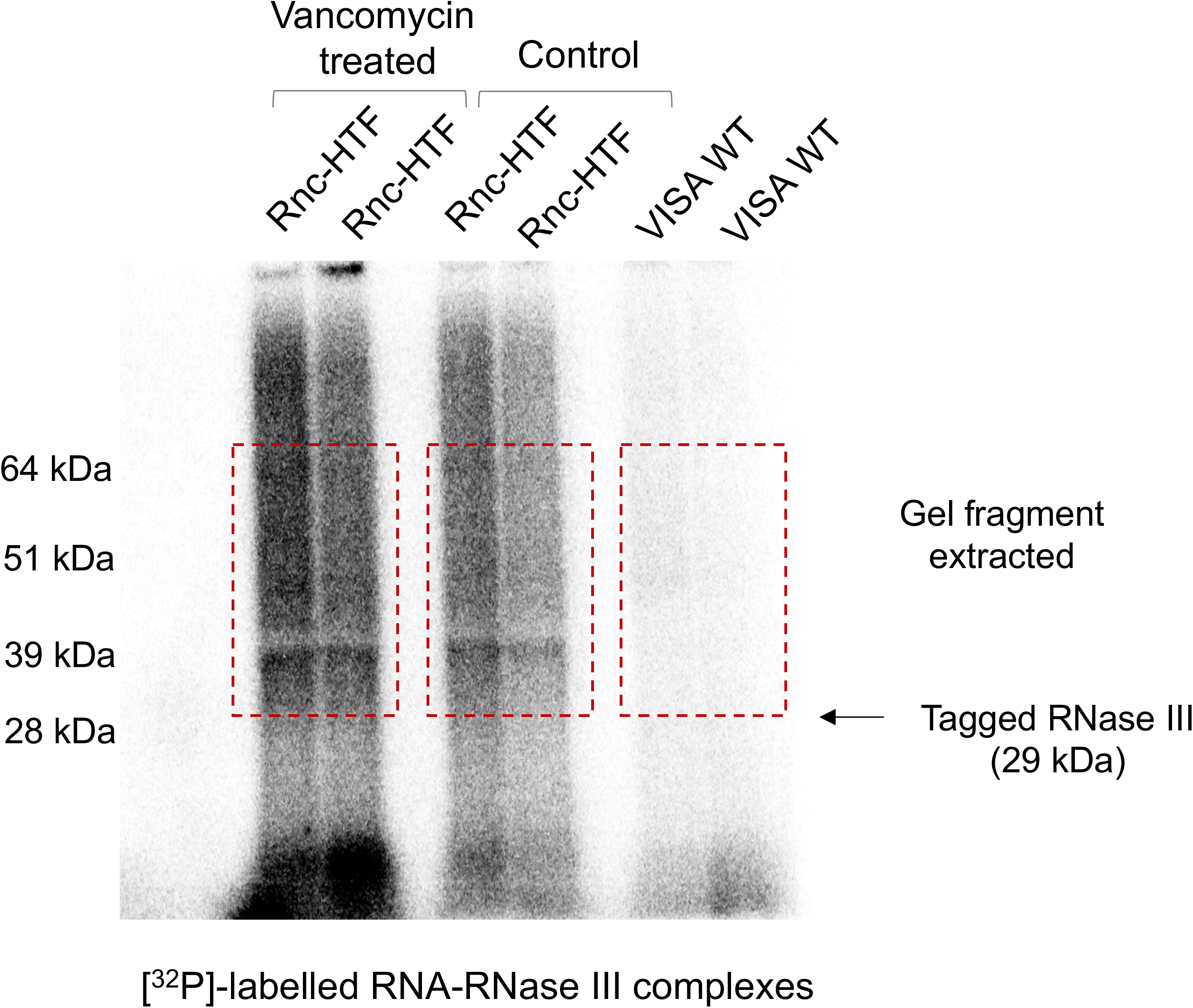
Autoradiogram of recovered RNA-RNase III complexes after affinity purification. Autoradiogram signal was observed after 2 hours of exposure and gel fragments from 28 kDa to 64 kDa outlined in the red box were extracted. Lane 1: vancomycin treated Rnc-HTF JKD6008 replicate 1. Lane 2: vancomycin treated Rnc-HTF JKD6008 replicate 2. Lane 3: untreated Rnc-HTF JKD6008 replicate 1. Lane 4: untreated Rnc-HTF JKD6008 replicate 2. Lane 5: wildtype JKD6008 replicate 1. Lane 6: wildtype JKD6008 replicate 2.

**Supplementary Figure 2.**
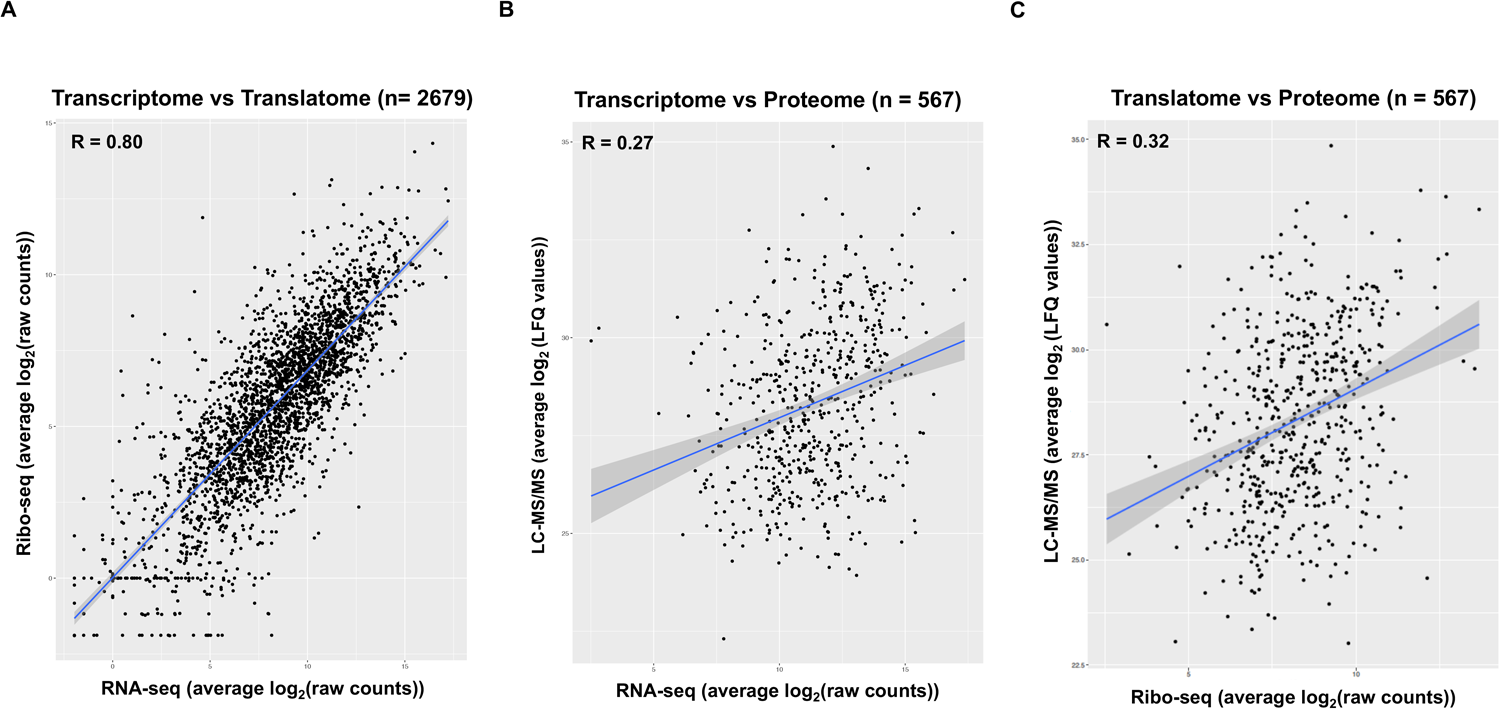
Correlations of gene expression levels between the RNAseq (transcriptome), Ribo-seq (translatome), and mass spectrometry (proteome) samples. **(A)** Average transcript abundance versus average ribosome occupancy. **(B)** Average transcript abundance versus average protein levels. **(C)** Average ribosome occupancy versus average protein levels.

**Supplementary Figure 3.**
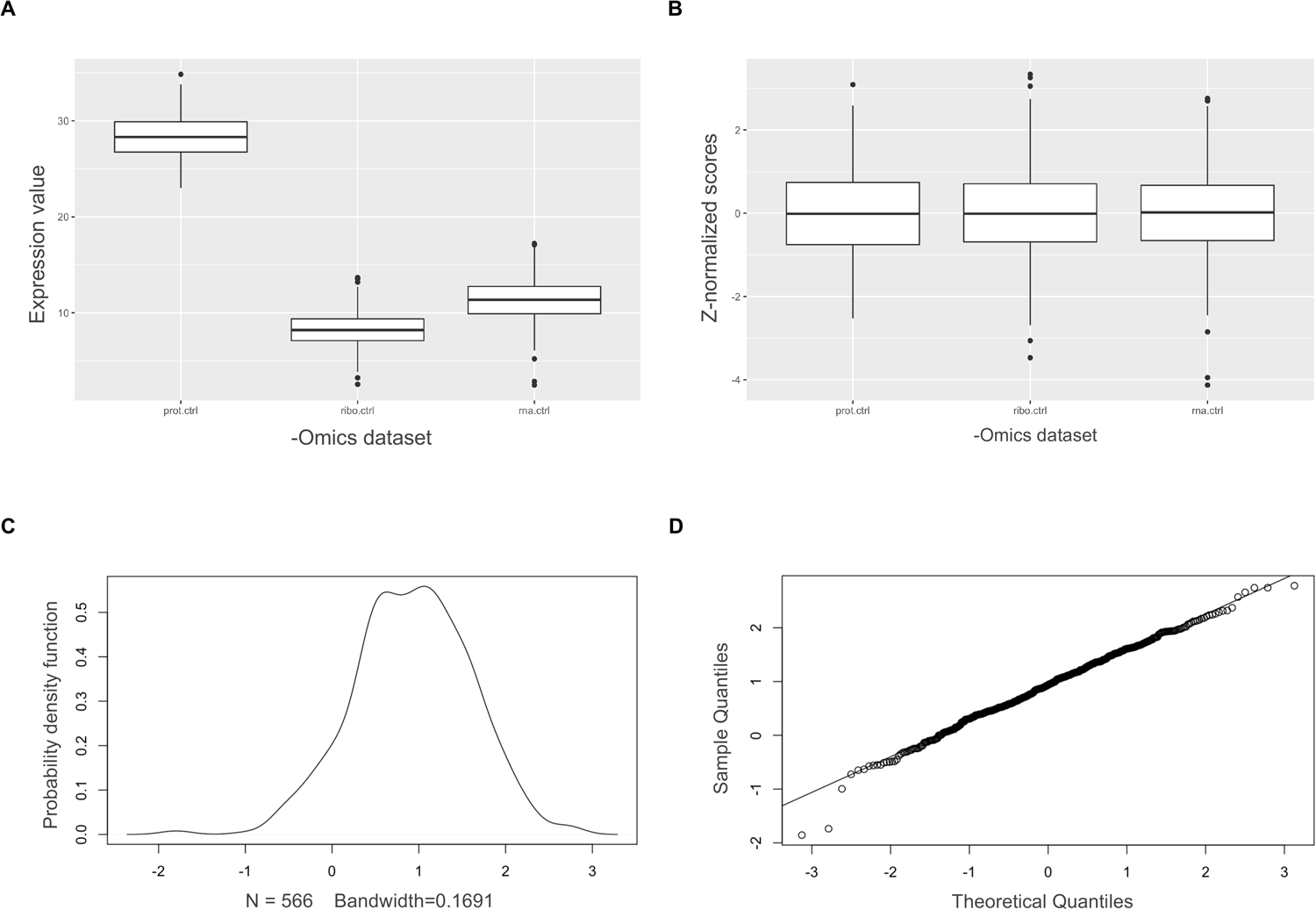
Quality control of the multi-omics datasets prior to SOMs clustering. (A) Boxplots of the average gene expression values for each -omics dataset prior to z-score normalisation. The expression values for the RNA-seq and Ribo-seq are represented as log_2_(raw counts) while the proteomics are represented as log_2_(label-free quantification value). (B) Boxplots of z-normalised values for each -omics condition. (C) Kernel-density plot and normal Q-Q plot (D) for the multi-omics datasets post z-normalisation.

**Supplementary Figure 4.**
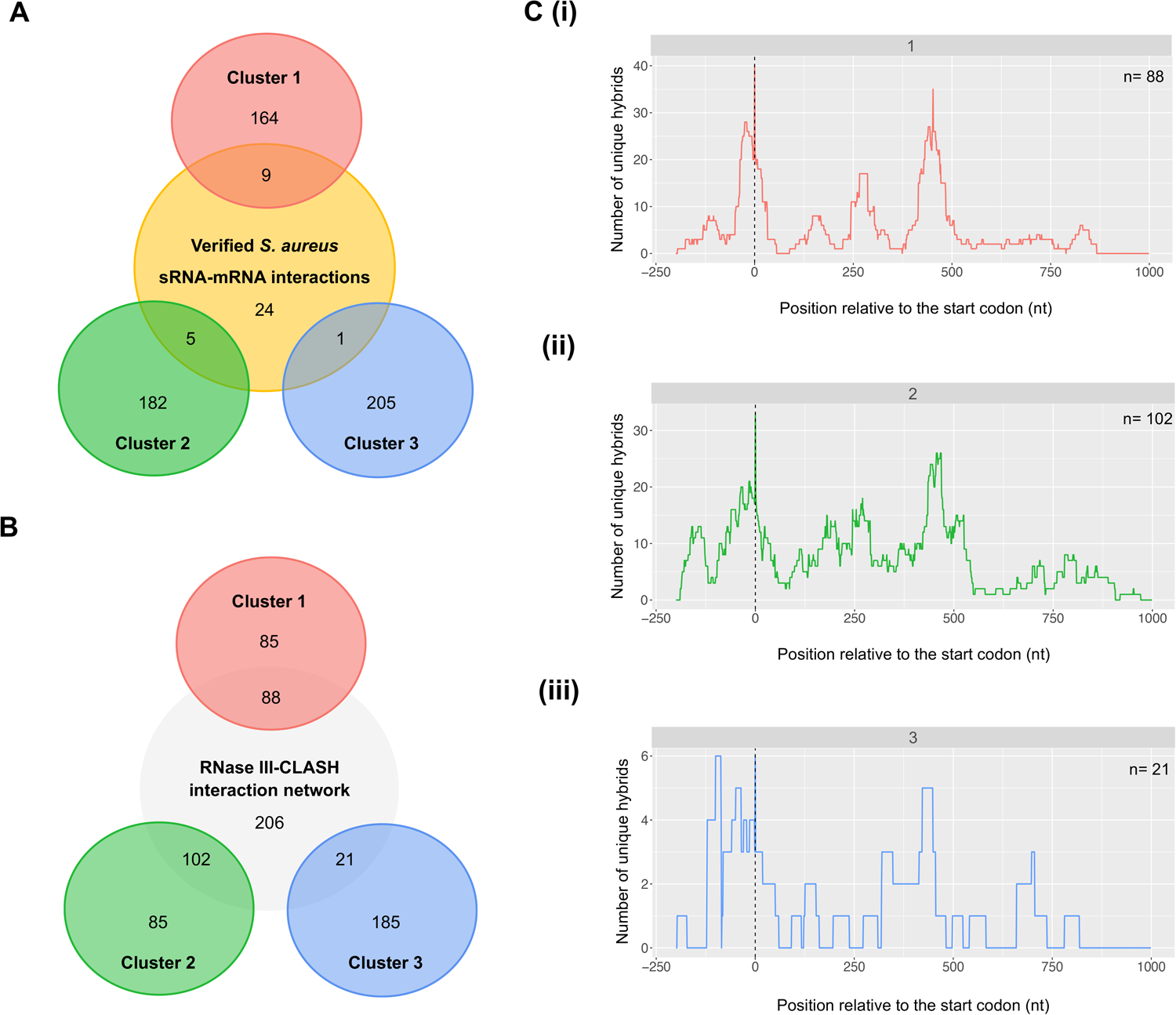
**(A)** Venn diagram of the 39 verified *S. aureus* sRNA-mRNA interactions overlayed with the SOMs clusters based on the mRNA targets. **(B)** Venn diagram of the 465 RNase-III CLASH captured sRNA-mRNA interactions overlayed with the SOMs clusters based on the mRNA targets. **(C)** A metagene plot was generated for sRNA-mRNA interactions that were detected in clusters 1 **(i)**, 2 **(ii)**, and 3 **(iii)**.

**Supplementary Figure 5.**
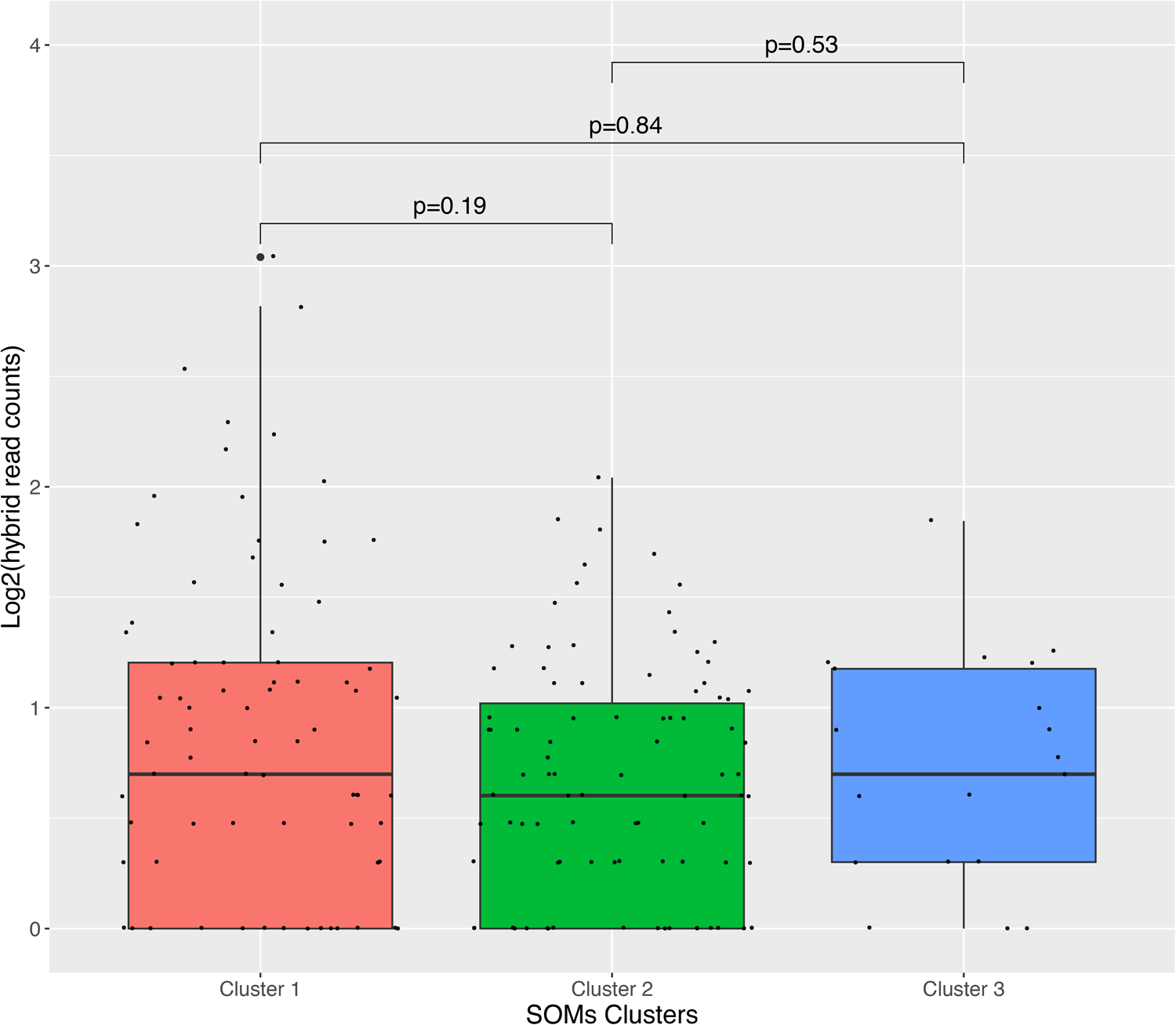
Distribution of sRNA-mRNA total hybrid counts in the 1×3 SOMs clusters. A boxplot was generated for sRNA-mRNA total hybrid counts that were detected in cluster 1, 2, and 3 (x-axis). The y-axis represents hybrid read count (log2).

**Supplementary Figure 6.**
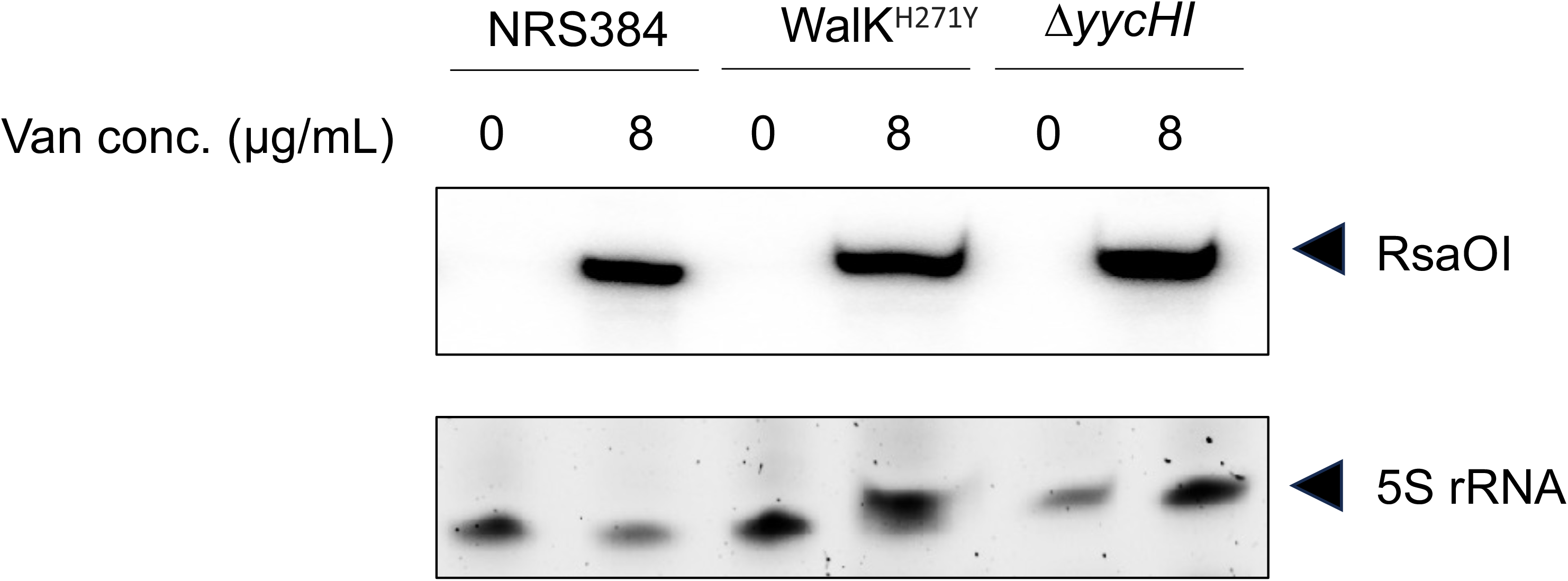
Northern blot analysis of RsaOI abundance in NRS384, WalK^H271Y^, and *ΔyyCHI*. Total RNA was extracted from the indicated strains treated with or without 8 µg/mL vancomycin for 30 minutes and then probed for RsaOI, All strains are derived from the USA300 strain NRS384.

**Supplementary Figure 7.**
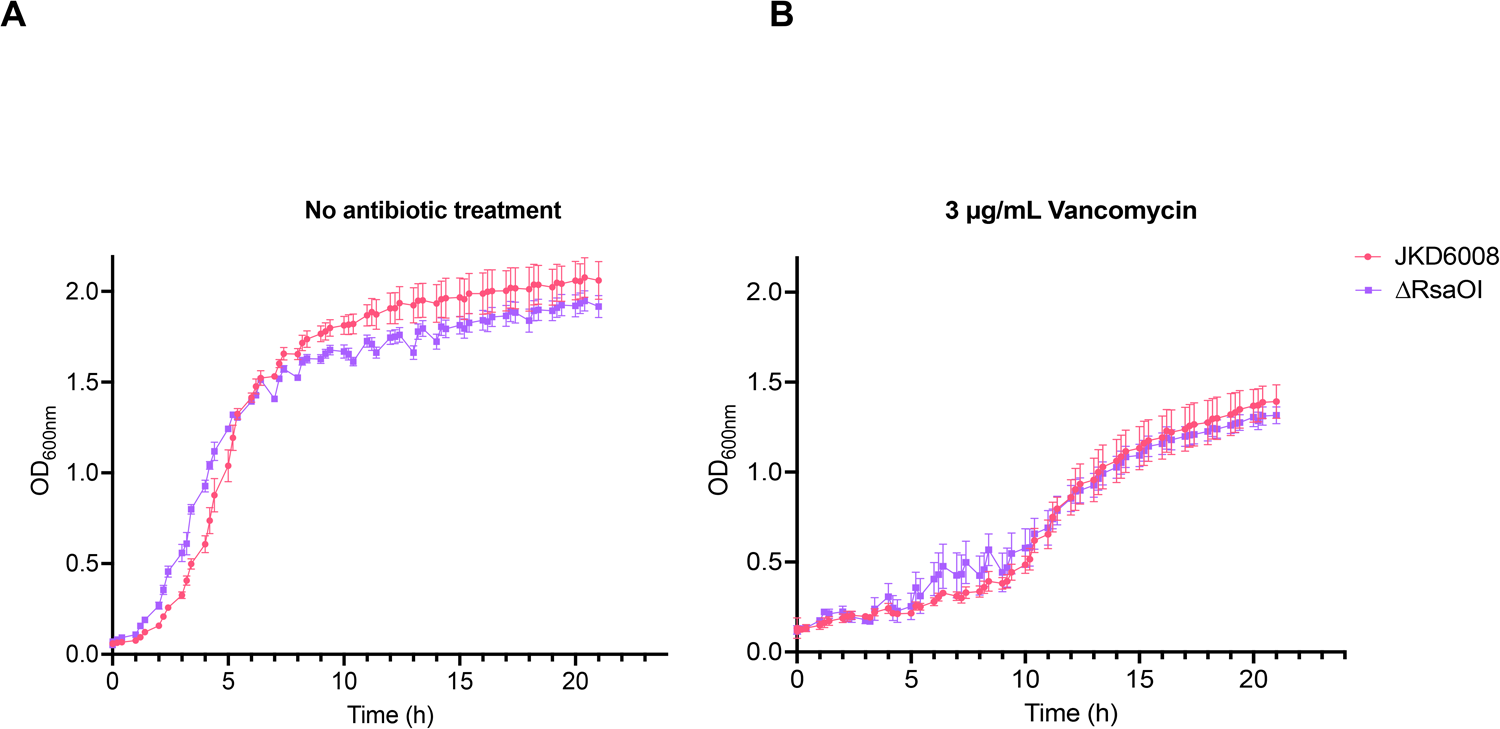
Growth curve analysis of VISA (JKD6008) and this isogenic *ΔrsaOI* mutant. Cultures were grown in MH media supplemented with or without 3 µg/mL vancomycin.

